# FIGL1 limits the formation of meiotic non crossover events in *Arabidopsis thaliana*

**DOI:** 10.1101/2024.09.30.615791

**Authors:** Delphine Charif, Raphaël Mercier, Joseph Tran, Christine Mézard

## Abstract

Meiotic recombination produces both crossover and non crossover events that are essential in the history of population genetics and evolution of species. In Arabidopsis thaliana, several pathways control the rate and distribution of crossovers. Here, by sequencing the four products of a series of tetrads, we confirm the antiCO role of *RECQ4A, RECQ4B, FIGL1* and *FANCM*. Moreover, when one of this gene is mutated, complex chimeric gene conversion events associated to crossover are observed suggesting a role of these proteins in limiting the multiple strand invasions. Nothing was known about the factors that could limit or increase NCOs. Here, we show that *FIGL1* plays a major role in controlling the NCO number.

## Introduction

Meiosis is central to sexual reproduction. During meiosis, four gametes are produced, each having a complementary haploid DNA content. In most species, recombination is essential for proper segregation of chromosomes during the first meiotic division. In addition, it generates diversity by reassorting parental alleles. Meiotic recombination is initiated by programmed double-stranded breaks (DSBs) in DNA ^1^. These breaks can be repaired using one of the three homologous templates present in meiocytes at this stage: the sister chromatid and the two non-sister chromatids of the homologous chromosome. When the DSB repair machinery uses one of the two non-sister homologous chromatids as a repair template, it leads to the formation of either crossover (CO), which is a reciprocal exchange between homologous chromosomes and thus gives a Mendelian segregation of the genetic markers, or non-crossover (NCO), which is the product of non-reciprocal exchange between homologous chromosomes, leading to a non-Mendelian segregation of genetic markers ^1^. If the DSB is repaired on the sister chromatid, it is genetically silent.

In most eukaryotes, the estimated number of programmed DSBs formed at the onset of meiosis significantly exceeds the CO number. This is particularly visible in plants, where tens of RAD51 foci that mark DSB repair sites are counted per chromosome, whereas the number of COs rarely exceeds three on a single chromosome ^2^. DSBs that do not form COs are repaired as NCOs or on sister chromatids, but the exact proportion of these two latter repair events is not known in most species.

COs can be formed by two different genetic pathways. One pathway is controlled by a group of proteins referred to as the “ZMMs”. ZMM dependent COs are more regularly spaced then if they occurred randomly along the chromosomes and they are called Class I or interfering COs. In most species, ZMM independent COs, or Class II COs, set a minor fraction (between 5 to 30%) of the total amount of COs in a single meiosis and their non-dependence on interference is under debate ^2,3^. However, it has been shown that in *Arabidopsis thaliana*, Class II CO’s number can be increased when either the FANCM complex ^4^, and/or the RECQ4A-RECQ4B-TOP3alpha-RMI1^5^ complex, and/or the FIGL1-FLIP^66,7^ complex are inactivated and in some combination of mutants their number can largely exceed the number of Class I COs (reviewed in ^2^).

Gene conversion (GC) events are a driver of gene and genome evolution. During meiosis, GC events include NCOs but can also be detected associated to COs hereafter called GCCOs. The number of NCOs formed in a single meiosis is difficult to estimate for a series of reason.

Formally, only events with a segregation 3 to 1 of a DNA sequence polymorphism in the four products formed after meiosis and without a reciprocal exchange between the flanking markers can be called NCOs ^8^. The recovery and analysis of the 4 meiotic products has been reported in a handful of species: some fungi including *Saccharomyces cerevisiae* ^9–11^, *Schizosaccharomyces pombe* ^12^ and *Neurospora crassa*^13^, mice^14^ and some plants like *Chlamydomonas reinhardtii* ^13^and *Arabidopsis thaliana*^13,15,16^. In species where the tetrad analysis is not available, the rate of NCOs has been estimated genome wide by whole genome sequencing of F2 or backcross off springs. Short events converting contiguous tracts of polymorphisms consistent with a non-reciprocal exchange of a DNA fragment between the two homologous chromosomes were considered as NCOs. Repair events that copy the homologous non-sister chromatid in a region that do not contain any polymorphisms are silent genetically. These “silent” NCOs could be largely underestimated if the average tract length is short and the density of polymorphisms is low. Moreover, very short events that convert a single SNP in regions that are difficult to sequence are also very difficult to detect and participate to the overall underestimate of the NCO rate. This explains why genome wide NCOs are easier to study in species with small genomes with a low complexity.

In this work, we analyze the pattern of meiotic recombination events *Arabidopsis thaliana* in tetrads from wild type and mutants of the Class II pathways. As previously reported, we show that in one single meiosis of *Arabidopsis thaliana* only a handful of DSBs are repaired as detectable NCOs ^13,15,16^. We also show that FIGL1 has a major effect in controlling the number of NCOs; when at least one of the CO pathways is inactivated in addition to increasing COs, complex recombination events are observed highlighting the role of these proteins in resolving complex recombination intermediates.

## Results

### Recovering the products of a single meiosis of A. thaliana

The process set up to obtain *A. thaliana* tetrad plants is detailed in the Method section. Briefly, to access to the full pattern of meiotic recombination events in a single meiosis, we crossed the two accessions Columbia and Landsberg *erecta* (Figure 1A). The two genetic backgrounds differ by around 0.4% at the nucleotide level ^17^. We selected for both accessions, a line having a mutation in the *QUARTET1* gene that results in the four pollen grains formed at the end of a single meiosis remaining attached in a tetrad ^18,19^. Each tetrad contains the whole history of the meiotic recombination events occurring in a single male meiosis. In order to investigate the recombination events of each pollen grain within a tetrad, a single tetrad was placed atop a solitary pistil of a Columbia plant. The pistil serves as the reproductive organ that contains female gametophytes. To prevent contamination caused by self-fertilization, we chose a Columbia accession with a mutation in the *ADENINE PHOSPHORIBOSYL TRANSFERASE 1* gene (*APT1, AT1G27450*), which results in male sterility (*apt1*-3)^20^. If every pollen grain from a single tetrad fertilizes an ovule successfully, the result will be fruits containing precisely four seeds. The seeds from these fruits were planted and grew into plants, from which DNA was extracted and analyzed with a series of markers positioned on the five chromosomes, differentiating between Columbia and Landsberg (SupTable1). To eliminate potential contamination from non-related pollen grains, only series of four plants displaying a Mendelian segregation of these markers were subjected to genome-wide DNA sequencing.

**Figure 1.**
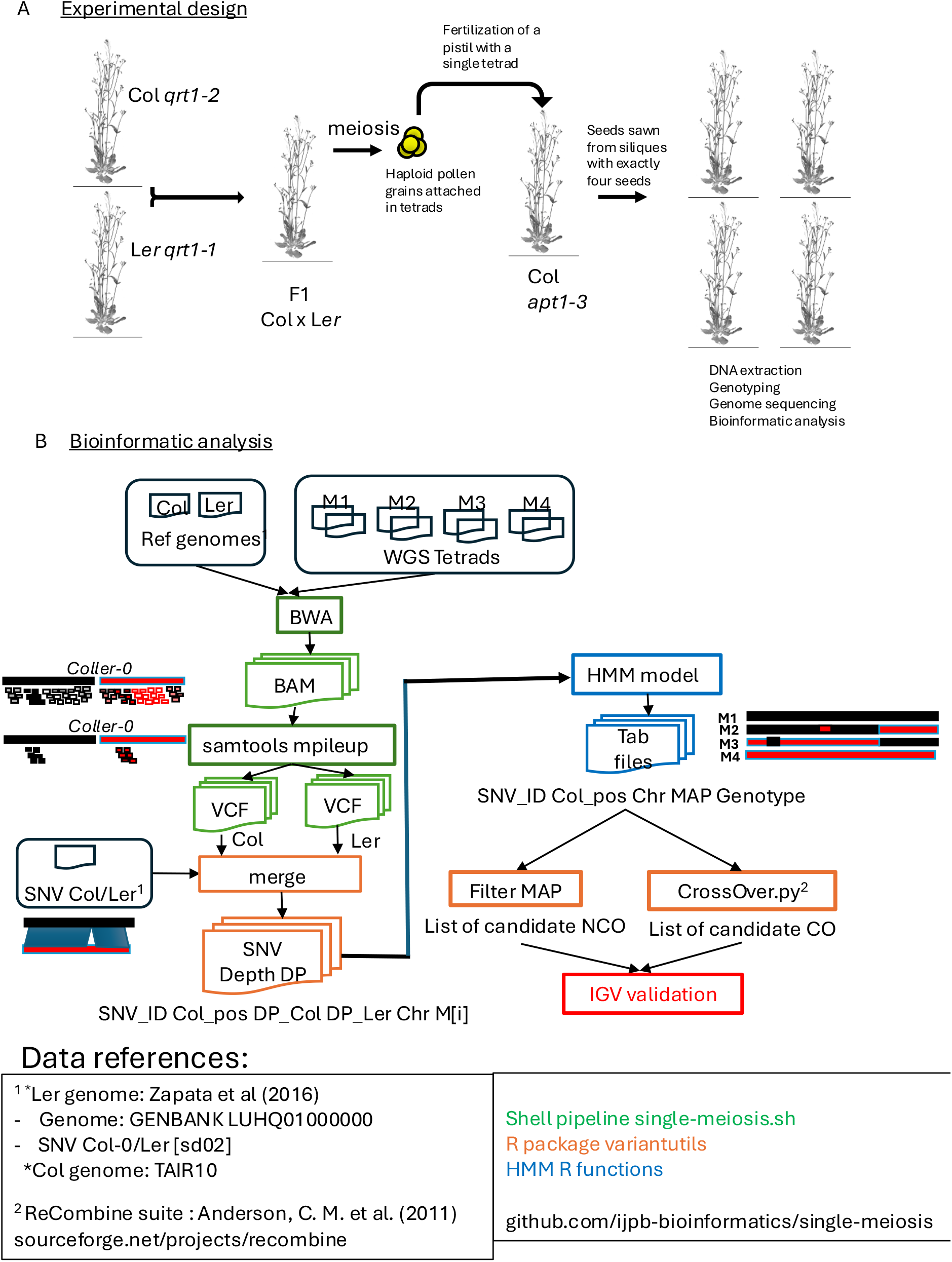
Experimental design. A. Schematic representation of the design to obtain tetrad of plants B. Schematic representation of the Bioinformatic pipeline

A total of 24 tetrads plus the two parental accessions Columbia and Landsberg *erecta* were sequenced (98 genomes). Among the sequenced tetrads 10 were wild-type and 15 contained mutation in one or a combination of genes that limit Class II CO formation: *fancm*^*-/-*^ (hereafter “Fa”; 2 tetrads), *figl1*^*-/-*^ (“Fi”; 2 tetrads), *recq4a*^*-/-*^ *recq4b*^*-/-*^ (“QAB”; 3 tetrads), *figl1*^*-/-*^ *fancm*^*-/-*^ (“FiFa”; 2 tetrads), *recq4a*^*-/-*^ *recq4b*^*-/-*^ *fancm*^*-/-*^ (“QFa”; 1 tetrad), *recq4a*^*-/-*^ *recq4b*^*-/-*^ *figl1*^*-/-*^ (“QFi”; 2 tetrads), *recq4a*^*-/-*^ *recq4b*^*-/-*^ *figl1*^*-/-*^ *fancm*^*-/-*^ (“QFiFa” ; 2 tetrads).

### Bioinformatic pipeline

The bioinformatic pipeline process set up to analyse the recombination events is detailed in the Supplemental data and summarized in Figure 1B. Briefly, we created a reference that contains the sequence of the 5 chromosomes from Columbia and the 5 chromosomes from Landsberg *erecta*. From the pool of SNPs/short indels that differentiate the two accessions Columbia and Landsberg published in a previous study ^17^, we retained a set of 522,658 that we could map unambiguously on the genome (hereafter referred as “Gold” SNPs/short indels)(available at **…**..). The mean distances between 2 consecutive markers were 43,50,48,45, and 43 for chromosomes 1,2,3,4, and 5 respectively) (SupTable2). Processed DNA reads from the tetrads were aligned on the reference without allowing mismatches and only reads that align on a single position, i.e. either on the Col or Ler, but not both, were kept. For each plant of the tetrad, the number of reads that mapped on each SNP/short indel were counted. This count was then used in a Hidden Markov Model (HMM) based pipeline to determine the most likely genotype Col or Heterozygous Col/Ler of each plant of a tetrad and also of the four plants of the tetrads at each given marker. For further analyses, COs were analyzed on all the tetrads, gene conversion events on all but 5 wild type tetrads because of a low sequencing coverage. Additionally, we had to remove one “QAB” tetrad of the NCOs analysis due to a low level of contamination of a DNA extracted from one plant of the tetrad with another DNA extracted from another plant of the same tetrad. Thanks to the marker genotypes obtained with the HMM model, COs were called and classified using the Recombine suite ^21^: COs are associated with SNPs with a mendelian segregation but with a change of continuity between the Columbia and Landsberg specific nucleotides along the DNA chromosomal molecule (cf SupFig1). Gene conversion events had a signature of a non mendelian segregation of one or a series of contiguous SNPs with one plant of the tetrad having one parental signature and the 3 others the corresponding SNPs of the other parents. The HMM model assigned a score from 0 to 20 (score= -log10(1-Posterior probability for the most likely state)) to both the gene conversion events associated to a CO and the gene conversion events not associated to COs so-called NCOs. We observed that the score was highly correlated to the tetrad sequencing depth (SupFig2). All the COs and the gene conversion events associated or not to COs detected with the pipeline were eye visualized in IGV to confirm or dismiss them whatever their score. Events dismissed were mostly due to very low coverage of the markers supposed to sign the events in one or several plants of the tetrads. We detected 475 COs on and 219 GCCO that were confirmed by eye visualization on IGV. 941 NCO events were detected by our pipeline with a score varying between 0.3 to the maximum 20. By eye visualization on IGV, we validated a set of 199 NCOs (SupTable3). We randomly picked up a series of 20 “eye” confirmed NCOs to be confirmed by Sanger sequencing. Their score varied between 0.3 and the maximum 20. 15 NCOs were fully confirmed. 4 NCOs could not be analyzed because the SNP(s) converted lied between highly repeated or indel regions between Columbia and Landsberg that precluded the analysis of the PCR product sequences. The last NCO, which had one of the lowest score among the series (0.36) was false negative: the markers supposed to be converted were not. We removed this NCO from the list as well as one other NCO that had a score that was lower to 0.36 leaving 197 NCOs for analysis.

### CO analyses

In the 10 wild-type tetrads, the number of COs varied from 6 to 14 with an average of 9.8 (+/-2.3)(Fig2A, SupFig1). 70% of the 98 COs were associated with a gene conversion events of a median length of 743 bp (minimum average length 256 bp-maximum average length 1,231 bp)(Fig2B, Table 1, SupTable4). In the mutant backgrounds, single mutation in *FIGL1* or in *FANCM* did not alter the CO number and when the 2 mutations were combined, just a slight increase of COs was observed (12 and 17 COs respectively in the two “FiFa” tetrads)(Fig2A). Inactivation of the two *RECQ4* (A and B) genes increased by around 2-fold the CO rates (24, 25 and 14 COs in the three “QAB” tetrads)(Fig2A). Inactivation of the two *RECQ4* genes associated to a mutation in either *FANCM, FIGL1* or both induced a significant increase of the CO number up to 62 COs in the “QFi” triple mutant (Fig2A). This huge increase of COs in a single tetrad led to a massive reassortment of the parental alleles in each single chromosome of the tetrad. While a single chromosome of a bivalent is rarely the recipient of more than 3 COs, the four chromosomes 5 of the “QFi” tetrad had 13, 12, 9 and 4 CO breakpoints respectively (SupFig1).

**Figure 2.**
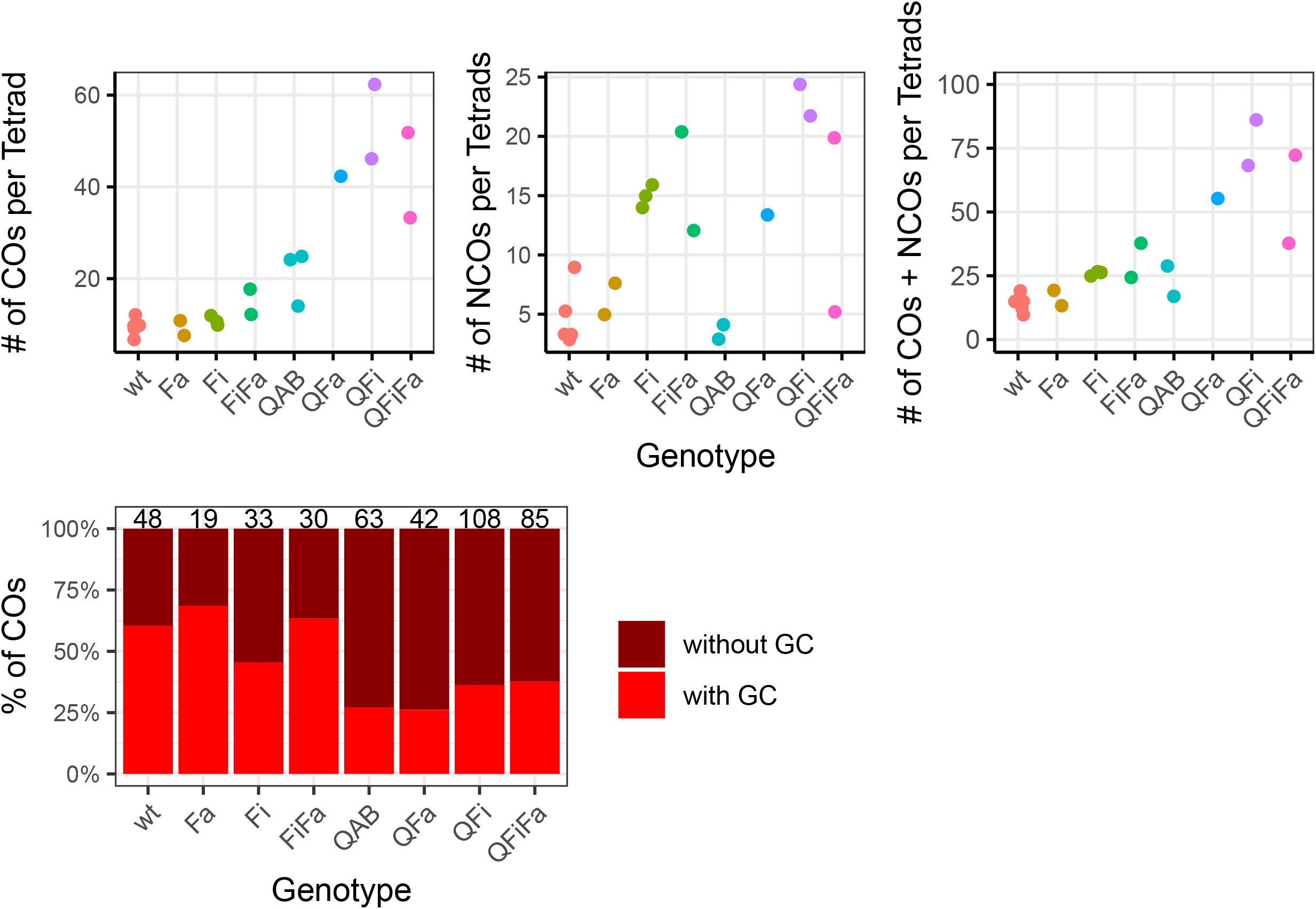
Count of the meiotic recombination events. A. CO and NCO numbers, B % of CO associated to a GC event

**Table 1.** Length of the GCCO and NCOs in bp.

|  | **wt** | **Fi** | **Fa** | **QAB** | **FiFa** | **QFa** | **QFi** | **QFiFa** |
| --- | --- | --- | --- | --- | --- | --- | --- | --- |
| n | 29 | 15 | 13 | 17 | 19 | 11 | 40 | 36 |
| GCCO midpoint length | 743 | 854 | 1019 | 979 | 1491 | 2146 | 1742 | 2061 |
| GCCO Minimum tract length | 256 | 430 | 271 | 306 | 721 | 1224 | 521 | 915 |
| GCCO maximum tract length | 1231 | 1609 | 1437 | 1654 | 2261 | 3068 | 2965 | 3208 |
| n | 23 | 45 | 13 | 7 | 32 | 13 | 46 | 25 |
| NCO midpoint length | 2019 | 1150 | 596 | 860 | 1114 | 3095 | 1370 | 1501 |
| NCO Minimum tract length | 185 | 2217 | 1166 | 1631 | 1951 | 5698 | 2526 | 2472 |
| NCO maximum tract length | 1853 | 83 | 26 | 89 | 271 | 492 | 216 | 532 |

### NCO analyses

We then looked at the NCO events (see Methods). In wild-type, we detected an average of 4.6 NCOs (+-2.6) per tetrads, which is half of the CO rate. Among the 23 NCOs, detected by our pipeline in the wild-type tetrads, 15 converted a single SNP. The minimum length between the 2 boarding non converted SNPs was 185 bp and the maximum length 3853 pb with a median length at 2019 bp (Table1). We estimated our probability of detection depending on the length of the NCO event (SupFig3). We randomly sorted 10,000 events with length varying between 1 to 10,000 bp and recorded the % of events that converted a SNP included in our Gold SNP set. When the events are shorter than 10bp the probability of their detection is less than 20% and the probability reached 87% when the tract length is longer than 7.5 kb (SupFig3). If NCOs had a tract length similar as the gene conversion events associated to CO (GCCOs) (743 bp, see above), 62% of the events would be probably detected. Thus our NCO rate is likely underestimated. In mutant tetrads, inactivation of *FANCM* or *RECQ4A* and *RECQ4B* does not modify the number of NCOs per meiosis. However, in *figl1* mutants, NCO increases by 3 to 4-fold . Combining a mutation in *FIGL1* with mutation in *FANCM* and/or *RECQ4A RECQ4B* does not further increase the NCO rate (Fig2A, SupTable3). We conclude that FIGL1 plays a major role in limiting the formation of NCOs.

When NCOs are added to COs, there are in average 14.2 events (+-3.2) in a single wild-type tetrad (Fig2A). A maximum of 84 recombination events were detected when *RECQ4A RECQ4B* and *FIGL1* were inactivated (Fig2A). Gene conversion events associated or not to COs modify the mendelian segregation of markers. In average, 24.8 SNPs were converted per meiosis giving a rate of 4.74×10^-05^ per base per meiosis. We noticed a slight increase of converted SNPs per meiosis when either *fancm* and/or *figl1* were mutated and it reached more than 150 when the whole series of *fancm figl1rec4a recq4b* was inactivated (SupFig4). No bias toward one or the other parent was observed.

### Complex gene conversion events

We “eye” visualized with IGV the 429 COs detected with the bioinformatic pipeline. When we analyzed the 32 GCCO detected in wild type tetrads, they were all simple with a single breakpoint on the two recombining chromatids (SupFig1, SupTable4). However, in mutant tetrads, 48 COs were “complex” with either several breakpoints occurring in the conversion tract and/or associated to a gene conversion events in a third chromatid not involved in the crossover events (SupFig5, SupTable4). These complex GCCOs were particularly detected when mutations in *RECQ4A RECQB, FIGL1* and *FANCM* were combined and could reach 72% of the events in the “QFa” tetrad. We did not observed events that involve the 4 chromatids. Among the NCO events, 2 in the *figl1* background and 1 in the double mutant *fancm figl1* exhibit discontinuous gene conversion events (SupFig5).

We also observed a long deletion of the first 309kb of the chromosome 5 in one plant of the tetrad RECQAB (see supFig6) which includes the first 180 genes of the chromosome. Thus, none of these genes is essential for pollen formation, pollen tube growth and fertilization even if some of them are preferentially expressed in the pollen.

### Localization of the recombination events

We examined the distribution of recombination events in various genomic regions such as promoters, 5’UTR, 3’UTR, introns, exons, transposable elements (TEs), and pericentromeres. The coding compartment was the largest, but it had the lowest density of “Gold” SNPs (Figure 3A-B). The highest density of “Gold” SNPs was found in the introns, promoters, and pericentromeres (Figure 3A). Recombination events are not localized proportionally to the size of the various compartments, neither to their “Gold” SNPs density (Figure 3A-B-C).

**Figure 3.**
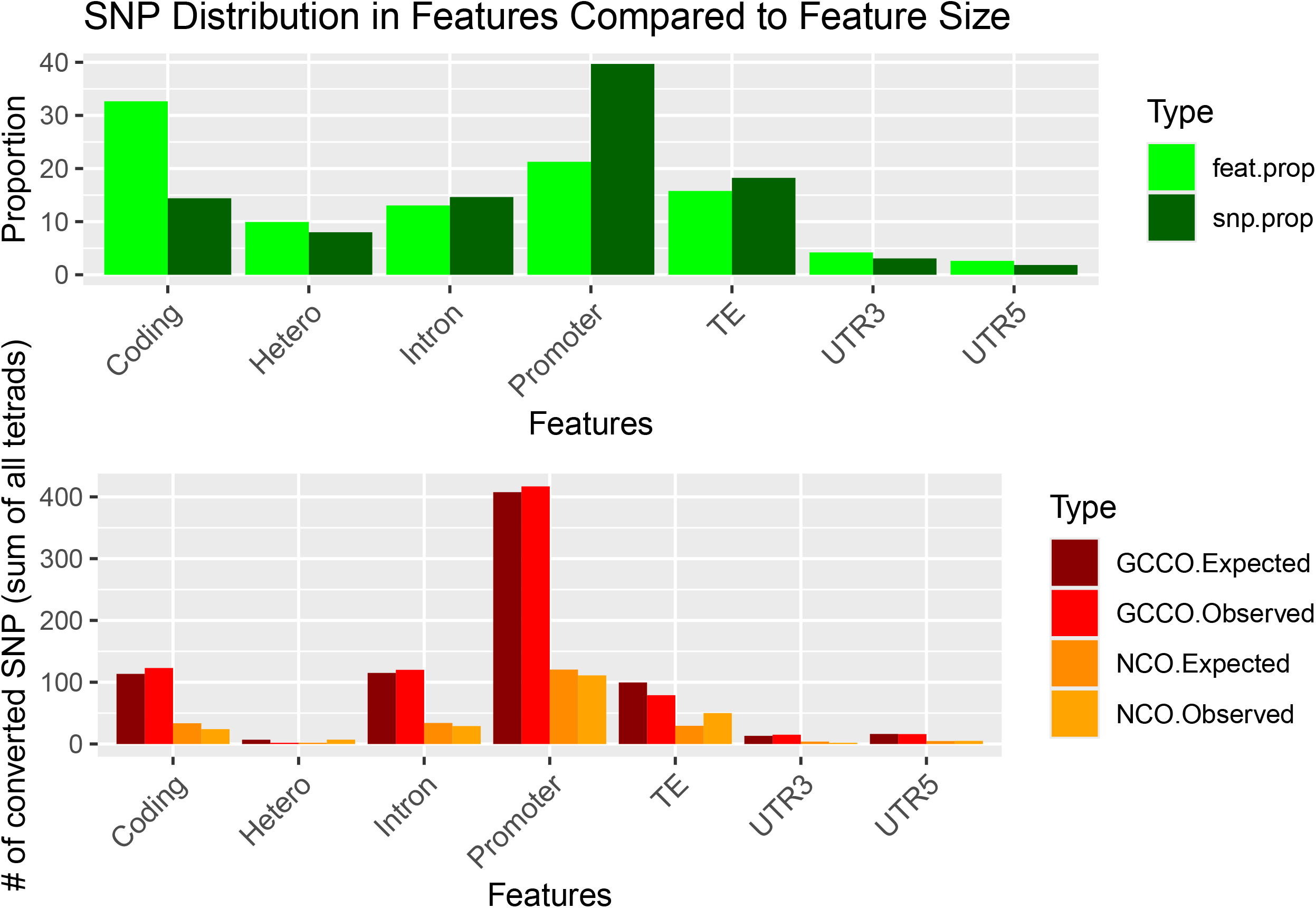
Localization of recombination events. A Relative size of the different genomic compartments relative to their SNPs density or their size in bp B Number of recombination events detected in the different genomic compartments relative to their expected number if the size of the different compartments is relative to their SNP density or their length in bp

Recombination occurs more frequently in introns and rarely in heterochromatin (11% and 8% of the size of the genome, respectively), whereas the promoters or coding regions (27 and 35% of the genome) have an observed number of recombination events proportional to their size (Figure 3C). When the SNP density is considered, recombination is found more frequently in promoters and less than expected in TEs (Figure 3B-C)). Thus, there are no clear features that influence the localization of meiotic recombination.

## Discussion

We analyzed of the recombinant genomes of a series of four *Arabidopsis thaliana* plants obtained by the fertilization of pistils with a single tetrad of pollen grains. The mother plants producing the pollen tetrads were F1 hybrids between the two accessions Col-0 and Ler, which differed by a set of reliable 522,658 SNPs. By comparing the DNA sequences of the four tetrad plants, the precise localization of all recombination events occurring in a single meiosis could be obtained and characterized.

As shown previously ^13,15,16^, COs and NCOs coexist during Arabidopsis meiosis. On average, 14.2 events (+-3.2) (9.8 (+/-2.3) COs and 4.6 NCOs (+-2.6)) were observed in a single meiosis. Our tetrads analysis confirmed that RECQ4A and RECQ4B play a major role in controlling CO formation, and when mutations in both helicases are combined with a mutation in either FANCM, FIGL1, or both, a 6 to 7 fold increase in the CO number per meiosis could be observed. However, for the first time, we analyzed the role of anti-CO pathways in the formation of NCOs. It has been hypothesized that the increase in COs observed in *recq4AB, fancm*, or *figl1* mutants was due to a redirection of the recombination intermediates supposed to be repaired as NCOs toward the class 2 COs pathway, leading to less NCOs and more COs ^22^. However, because of an already very low level of NCOs in wild-type meiosis, our analysis does not have the power to detect a decrease in the NCOs rate. Nevertheless, mutating *FIGL1* either alone or in combination with *fancm* or *recq4ab* had the opposite effect with a 3 to 5 fold increase. As suggested previously in ^23^, our results suggest that FIGL1 may play a role in the strand invasion step. We also observed complex GC-CO events in all mutant backgrounds except for the single *fancm* mutation. These complex GC-CO events were either discontinuous tracts or GC events in a chromatid not involved in the CO event, which is compatible with multiple invasion steps ^24^ in the absence of these proteins. GC tracts have a median length at least twice as long when formed by an NCO than when associated with a CO which is different from what has been observed in *S. cerevisiae* where GC-COs are slightly longer than NCOs. This apparent difference could be due to a lower SNP density which does not allow for the identification of all NCO events and/or very short polymerase tracts. By counting RAD51 or DMC1 foci on meiotic chromosome spreads, the DSBs number per male meiocyte has been estimated to be approximately 200 in *A. thaliana* ^25^. Only a small fraction of these DSBs (9.8 on average in our study) are channelled toward the CO pathways. Several analysis including this work have shown that when anti CO pathways are inactivated, a hundred of recombination intermediates are converted into COs ^22,26,27^. In this study, we report that in a *figl1* mutant, we observe a 5 fold increase in the NCO rate, with a maximum of 22 events in a single tetrad which is likely underestimated due to the low density of SNPs (SupFig3). If we combine these two results, in a single meiosis, half of the DSBs could be repaired as either COs or NCOs; however, in wild-type meiosis, several pathways limit the formation of recombinant products. Why do so many DSBs are made if only a small fraction is converted into COs ? A large fraction of DSBs could be chanelled to invade homologous non-sister chromatids. Most of these invasions would not activate a DNA polymerase to copy the template or if so, the polymerase would copy just a very short patch that would not lead to a detectable gene conversion event in most cases. However, these invasion events distributed all along the chromosomes would drive the proper alignment of homologous chromosomes.

## Material and Methods

### Plant materials and growth conditions

*A. thaliana* plants were grown in greenhouses with 70% humidity and under a 16 h/8 h day/night photoperiod with temperatures of 19 °C during the day and 16 °C at night. Wild-type Col-0 and Ler-1 are 186AV1B4 and 213AV1B1 from the Versailles *A. thaliana* stock centre (http://publiclines.versailles.inra.fr/).The following mutations were used in this study: in Col-0, *fancm-1* ^4^, *figl1-1*^6^, *recq4a-4* (N419423) and *recq4b-2* (N511130) ^28^, *apt1*-3 ^20^, *qrt1-2* (CS8846)^18^; in Ler-1, *fancm-10* ^6^, *figl1-12* ^6^, *recq4a-W387X* ^5^, and *qrt1-1* (CS8845)^18^. Flowers from the F1 obtained by crossing CS8846 and CS8845 were tapped on a glass slide to deposit tetrads of pollen grains. Using a needle, a single tetrad was taken and gently embedded in the stigma of a *apt1*-3 Col-0 plant. The fertilized plant was grown in a greenhouse without other Arabidopsis plants to avoid cross-pollinization. Seeds from fruits having exactly four seeds were sterilized a few seconds in a drop of 70% ethanol and sown on solid medium ^29^ without sugar. After stratification for 2 days at 4°C, plants were grown in long day conditions. At the four leaves stage, the plants were transferred on soil and grown in green houses until the induction of the first hamp.

### Methods

DNA was extracted from the set of four plants as described in and analyzed running a set of ten specific PCRs that allow to identify specific Landsberg/Columbia markers along the five chromosomes (SupTable1). Only the series of 4 plants that show a mendelian segregation of these ten markers were further analyzed. Leaf samples were used for DNA purification and library preparation for Illumina sequencing (HiSeq 3000 2×150pb) performed either by the GeT-Plage center, France (https://get.genotoul.fr) or the EPGV center, Evry, France (https://www.genopole.fr/genopolitains/laboratoires/epgv/).

A total of 24 tetrads were sequenced plus the parent Columbia and Landsberg *erecta*. For each tetrad, the mutant genotype for *RECQ4A, RECQ4B, FIGL1, FANCM* was confirmed on IGV.

## Data availability

Sequencinq data are accessible under https://www.ebi.ac.uk/biostudies/arrayexpress/studies/E-MTAB-14435

## Supplementary

The Bioinformatic pipeline, statistic codes, supplementary methods, supplementary figures and supplementary tables are available under https://github.com/ijpb-bioinformatics/single-meiosis/

## Aknowledgments

We are indebt to Stéphane Robin for producing the HMM model to obtain the gene conversion events. Aurélie Chambon, Laurène Giraut, Jan Drouaud, Vanessa Zanni, Mathilde Séguéla, Joiselle Fernandez, Cécile Larchevèque, Sarah Bellalou produced the genetic material. Olivier Martin, Matthieu Falque, Matthias Zytnicki were involved to produce the bioinformatic pipeline. Mathilde Grelon was involved in the writing of the paper. We are grateful to the genotoul bioinformatics platform Toulouse Occitanie (Bioinfo Genotoul, https://doi.org/10.15454/1.5572369328961167E12) for providing help and/or computing and/or storage resources.

We would like to thank Peter Schlögelhofer for entrusting us with 5 wt tetrads to analyze and for his support on this project.

